# Visuo-Vestibular Integration for Self-Motion: Human Cortical Area V6 Prefers Forward and Congruent Stimuli

**DOI:** 10.1101/2024.11.27.625640

**Authors:** Sarah Marchand, Marine Balcou, Philippine Picher, Maxime Rosito, Damien Mateo, Nathalie Vayssiere, Jean-Baptiste Durand, Alexandra Severac Cauquil

**Author notes:** Alexandra Séverac Cauquil, CerCo Centre de Recherche Cerveau et Cognition (CNRS/Université Toulouse III), Pavillon Baudot-CHU Purpan, Place Baylac 31300 Toulouse cedex – France.

## Abstract

**BACKGROUND:** The integration of visual and vestibular input is crucial for self-motion. Information from both sensory systems merges early in the central nervous system. Among the numerous cortical areas involved in processing this information, some (V6 and VIP) respond specifically to vestibular anteroposterior information.

**OBJECTIVE:** To further understand the involvement of these and other areas in self-motion processing when vestibular and visual information are combined with varying congruence and direction parameters.

**METHODS:** Fifteen subjects underwent an MRI session while receiving visual (optic flow patterns) and galvanic vestibular stimuli mimicking six conditions: (1) visual forward, (2) visual backward, visual forward with (3) congruent or (4) incongruent vestibular information, visual backward with (5) congruent or (6) incongruent vestibular information.

**RESULTS:** The combination of concurrent vestibular stimulation and fully consistant optic flow patterns activated several bilateral cortical areas found predominantly in the insula. Among these vestibular areas and those previously defined in our initial study, the large majority do not show any specifity for the forward/backward direction or for the visuo-vestibular congruency. A notable exception was the parieto-occipital area V6, which showed a marked preference for congruent visuo-vestibular signals and for cues signaling forward motion.

**CONCLUSIONS:** By showing that V6 is more active when visuo-vestibular signals are more ecological (*i.e.* when both signals specify the most common self-motion direction), our results support the view that this area plays a crucial role in visuo-vestibular integration during self-motion.

## Introduction

Forming robust and dynamic representions of our body in space is essential for the control of posture and self-motion. These representations are produced by integrating signals from different modalities, notably of visual, vestibular and muscular proprioceptive origins [5, 12, 50, 52]. Studies have shown that altered visuo-vestibular integration disturbs self-motion [2, 7, 37, 43, 44, 64] and can lead to symptoms of nausea, vertigo or dizziness [13], as is the case in Persistent Perceptual Postural Dizziness or PPPD [62].

Electrophysiological studies in non-human primates have identified several cortical areas in which neurons respond to both visual and vestibular self-motion signals, particularly in the temporal lobe, around both the temporoparietal junction and the midposterior Sylvian fissure. These regions notably include the medial superior temporal area (MSTd) [27], the ventral intraparietal area (VIP) [21], the visual posterior sylvian area (VPS) [20, 41], the frontal eye field area (FEF) [40] and region 7a [6]. Correspondingly in humans, functional imaging studies have revealed that the network of cortical areas responding to both visual and vestibular self-motion signals is extensive [60], encompassing hMSTd [28], hVIP [10] and CSv [59]. Overall, both the regions involved [61, 22] and their connectivity pattern [23, 59] seem to be highly conserved between these primate species. A notable difference is in area V6, which responds more strongly when optic flow patterns are consistent (rather than inconsistent) with self-motion in humans [61] but not in monkeys [22].

Interestingly, by using galvanic vestibular stimulation (GVS) [26, 51, 59] and functional MRI in humans, we recently demonstrated that among the visuo-vestibular regions identified, only two —V6 and VIP areas— respond to GVS when it specifies a vection along the anteroposterior axis, but not when it mimicks a lateral vection [1].

In the present study, we further investigate these observations by combining visual and vestibular signals that can independently specify forward and backward directions along the anteroposterior axis. Our aim is to uncover whether some of the involved regions are more specifically involved in visuo-vestibular integration during ecological conditions of self-motion (*i.e.* when visual and vestibular signals congruently specify forward locomotion).

## Material and methods

### Participants

Subjects were recruited via posters and provided with information about the research, a consent form and a minimum 5-day withdrawal period. Fifteen healthy participants were included in the present study (mean age: 23 years, range: 21-26 years, 10 females, 5 males). Fifteen healthy participants were included in the present study (mean age: 23 years, range: 21-26 years, 10 females, 5 males). All subjects had normal or corrected-to-normal visual acuity. To be recruited, the subjects were required to have (1) no contraindication to MRI, (2) no history of vertigo nor motion sickness, (3) no clinically significant conditions preventing them from staying in the scanner for 1 hour in the scanner without discomfort, (4) no progressive psychiatric or neurological pathology, (5) a sufficient proficiency in French to understand the consent form and (6) no suspicion of pregnancy. This study was approved by an ethics committee (ID CPP 15-001/2014-A01893-44). All subjects gave their written informed consent before their participation and received 60 euros of monetary compensation.

### Visual stimuli

The visual stimuli were projected by an overhead projector located outside the scanner room onto a screen at the entrance of the MRI scanner tunnel. A mirror located above the MRI antenna and directed towards the screen at the entrance to the tunnel allowed subjects to see the screen while lying in the scanner. The viewing distance was 82 cm and the diagonal of the monitor was 40 cm long. The stimuli consisted of 2 seconds of optic flow videos composed of white random dots moving against a black background (Fig. 1A). This short stimulation duration was chosen to avoid motion after-effects and minimize potential postural responses while being sufficient to trigger robust brain activations [1], notably in area V6 [16]. The videos were displayed at a resolution of 800 x 600 pixels, covering 30 x 23 degrees of visual angle, with a refresh rate of 85 Hz and a motion-coherence level set at 30% (which means largely unambiguous optic flow patterns). The dots were either expanding to mimic forward self-motion, or contracting to mimic backward self-motion.

**Fig. 1.**
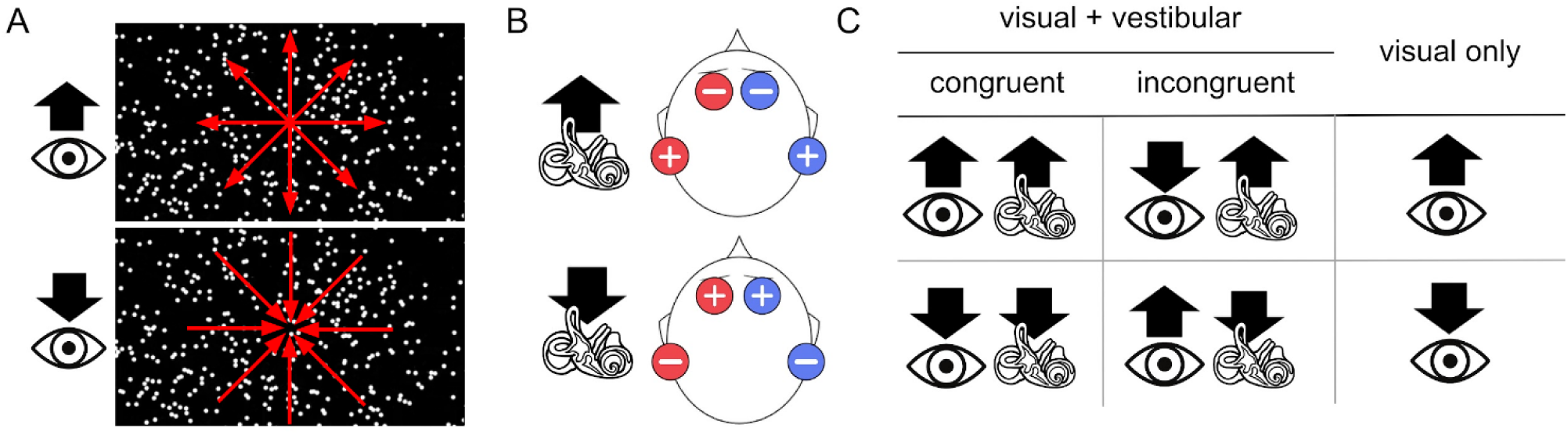
Study paradigm. A/ Visual stimuli. Image captures from the optic flow videos presented to the subjects (superimposed red arrows indicate the expanding or contracting movement of the dots). The large black arrows above the eye logo represent the direction of the visual self-motion illusion (forward for expanding dots and backward for contracting dots). B/ Vestibular stimuli. The two pairs of electrodes, with one anode (+) and one cathode (-) each, are designated by their red and blue colors, respectively. The large black arrow above the vestibule logo indicates the direction of the vestibular self-motion illusion (forward or backward). C/ Conditions. The black arrows represent the direction of the specified self-motion direction. The eye represents optic flow stimulation and the vestibule represents GVS stimulation. The experimental design includes six conditions. Two conditions (first column) consisted of multimodal (visual + vestibular) stimulation with congruence in the direction of the self-motion illusion (either both forward or both backward). Two conditions (second column) were also multimodal, but with the two modalities being incongruent (one induced a forward self-motion illusion, the other a backward self-motion illusion). Finally, two conditions (third column) involved only the visual modality, inducing an illusion of either forward or backward self-motion. During the acquisition, these six conditions were randomly distributed in 60 trials for each run (10 trials per condition) and 8 runs were acquired per subject.

### Galvanic vestibular stimulation

Before positioning the electrodes, the subject’s skin was prepared with alcohol and the electrodes were covered with conduction gel. As shown in Fig. 1B, the position of the 4 carbon MRI electrodes (Skintact, FSWB00) followed a monopolar binaural setup, with two configurations: 1/ to mimic an input of forward motion: the anodes (+) of each of the two pairs of electrodes were placed behind the ear, at the level of the mastoid processes and the cathodes (-) were placed ipsilaterally, on the subject’s forehead, just above the eyebrows 2/ to mimic an input of backward motion: anodes (+) of each of the two pairs of electrodes were placed at the level of the forehead and the cathodes (-) were placed ipsilaterally, at the level of the mastoid processes [57, 58]. Two stimulators (Digitimer DS5 CE certified for biomedical research N(IEC) 60601) located outside the scanner room, one for each pair of electrodes, delivered the electrical current via MRI compatible cables running through a waveguide. The vestibular stimulation consisted of 2 seconds of square current of 1 mA intensity between each mastoid process and the subject’s forehead. As with visual stimulation, the stimulation time and intensity were lower than in other studies (*e.g.* [54]) to minimize the risks of postural responses, current diffusion and discomfort while being sufficient for triggering robust cortical activations [1]. Subjects reported that neither pain nor discomfort were experienced during the stimulation, which enabled us to avoid the use of potentially allergenic local anesthetics.

### Design

Data were collected using an event-related design. Each run lasted 7.5 minutes. The six defined combinations resulting from the pairing of these two stimulation modalities enabled the creation of two “Congruent” conditions (in which the visual and vestibular organs are simultaneously stimulated by input in the same direction, either forward or backward), two “Incongruent” conditions (in which the visual and vestibular organs are stimulated by input in opposite directions, one forward, the other backward) and two conditions in which only the visual stimulation was delivered (Fig. 1C). In all conditions, the stimulation, whether visual or multimodal, lasted 2 seconds, with an inter-stimulus interval varying randomly between 5 and 7.5 seconds. The conditions were applied in a pseudo-random order and 8 runs of 60 simulations were performed per subject.

### Data acquisition

The MRI images were acquired by a clinical 3T scanner (Philips Achieva), equipped with a 32-channel head coil. At the beginning of each session, a high-resolution T1-3D anatomical image (4 minutes) was acquired (240 slices; repetition time (TR): 7.4 ms; echo time (TE): 3.4 ms; flip angle (FA): 8, percent phase FOV: 100; voxel size 1 x 1 x 1 mm). Functional EPI images for localization of congruent and incongruent visual and vestibular stimuli processing (1 hour) were acquired in the form of 8 runs consisting of 10 blocks of 6 stimuli according to an event-related design, with a TR of 2.5 seconds, a TE of 30 ms, a voxel size of 2 mm, a slice thickness of 3 mm (gap between slices = 0 mm), an FA of 90 degrees, a SENSE factor of 2.8 and an EPI factor of 39. Each run comprised 184 volumes of transversely oriented slices that covered the whole brain.

### Data analysis

#### Pre-processing

The functional volumes underwent a conventional pre-processing pipeline, starting with spatial and temporal realignments with the SPM12’s dedicated tools in MATLAB. Volume and slice outliers were detected and repaired using SPM12’s ArtRepair module [48]. The realigned and corrected images were then coregistered to the anatomical image acquired during the same session. The anatomical image then underwent segmentation and normalization in MNI space, and the normalization parameters were finally applied to the functional volumes before spatial smoothing (gaussian kernel, FWHM = 4 mm isotropic). All of the following statistical tests were carried out using the MATLAB interface with the Statistics and Machine Learning toolboxes.

#### Whole brain analysis

For each subject, we performed a 1^st^ level whole-brain analysis using a general linear model (GLM) implemented in MATLAB/SPM12, and then we carried out a 2^nd^ level group analysis for generalization. We mapped the cortical regions whose activation increased when simultaneous vestibular and visual information was provided by contrasting all the “Visual + vestibular” conditions (the 4 conditions in which both modalities were stimulated) with the “Visual only” conditions (the 2 conditions in which only the visual modality was stimulated). A results threshold was set at p < 10^-3^ (uncorrected) with a minimum cluster size of 5 voxels. The main activation clusters of the network were labeled using the Anatomy3 toolbox with probabilistic cytoarchitectonic maps in the MATLAB/SPM12 interface [30], the Brainnetome atlas [33] and comparison with similar coordinates in the literature [35, 36, 42, 46, 67] (Table 1). To visualize this functional network on a cortical surface, we used Caret5 software [65].

**Table 1.** Coordinates of the main activation clusters (local maxima) from the “Visual + vestibular” > “Visual only” t-contrast in the whole-brain second level analysis.

| # | label | anatomy | left coordinates |  |  |  | right coordinates |  |  |  |
| --- | --- | --- | --- | --- | --- | --- | --- | --- | --- | --- |
|  |  |  | X | y | z | t-value | x | y | z | t-value |
| 1 | AI | ventral granular insula | -34 | 4 | -12 | 8.64 | 38 | 6 | -6 | 9.03 |
| 2 | OP3 | dorsal granular insula | -38 | -6 | 4 | 9.15 | 40 | -2 | 6 | 6.15 |
| 3 | OP8 | frontal opercular cortex | -32 | 10 | 12 | 10.45 | 34 | 8 | 14 | 4.24 |
| 4 | PIVC | dorsal granular insula | -40 | -10 | 12 | 8.16 | 36 | -10 | 16 | 6.59 |
| 5 | OP4 | central opercular cortex | -56 | 2 | 8 | 6.67 | 54 | -10 | 14 | 7.22 |
| 6 | PIC | postcentral gyrus | -62 | -16 | 20 | 6.63 | 60 | -22 | 22 | 6.93 |
| 7 | PFcm | planum temporale | -62 | -38 | 16 | 5.89 | 52 | -32 | 14 | 4.82 |

#### Region of interest (ROI) analysis

To investigate which of these regions were sensitive to the simultaneous solicitation of vestibular and visual modalities sollicitation (considered here as a positive control), to visuo-vestibular congruence, and to the direction of visuo-vestibular input, we used a region of interest (ROI) approach. This analysis was carried out using the coordinates of the regions extracted from the whole brain analysis on the “Visual + vestibular” > “Visual alone” contrast. To avoid statistical circularity, the data set was split in half. ROIs were determined by using only the even-numbered runs for all 15 subjects and their sensitivity profiles were evaluated on the odd-numbered runs only. We generated ROIs in the form of spheres of 5 mm radius (representing 65 voxels) from each set of coordinates using SPM12’s MarsBaR toolbox [11]. Once the ROIs were obtained, we extracted, for each subject and for each ROI, the mean contrast value for the “Visual + vestibular” > “Visual only” contrast (as a positive control), “Congruent” > “Incongruent” contrast and “Forward” > “Backward” contrast. The “Congruent” > “Incongruent” t-contrast images were obtained by comparing the activations of the congruent conditions (“Visual + vestibular forward” and “Visual + vestibular backward”) with those of the incongruent conditions (“Visual backward + vestibular forward” and “Visual forward + vestibular backward”). The “Forward” > “Backward” t-contrast images were obtained by comparing the activations elicited by forward conditions (“Visual only forward” and “Visual + vestibular forward”) with those of the backward conditions (“Visual only backward” and “Visual + vestibular backward”). Once the mean contrast values had been obtained, we performed t-tests (one sample t-test, one-tail, hypothesized mean contrast value > 0, significance threshold: p < 0.05) on the mean contrast values of the images from our contrasts of interest for each ROI, to determine whether the activation of some of these regions was significantly different between the two conditions of our different contrasts.

In order to determine the effect of our different conditions on regions known to be activated by GVS in the absence of visual stimulation, we conducted a second ROI analysis with the coordinates taken from our previous study (Table 1) [1]. These coordinates, initially provided in Talairach space, were converted to MNI space using the free online resource MNI ˂-˃ Talairach Tool (BioImage Suite, Yale School of Medicine) [49] (Table 2). We generated spherical ROIs with a radius of 5 mm from these MNI coordinates using MarsBar [11] and we extracted the values from our contrast images in the spheres constituting the ROIs. We used the same contrasts of interest: “Visual + vestibular” > “Visual only”, “Congruent” > “Incongruent” and “Forward” > “Backward” to define the effect of the simultaneous visuo-vestibular information, the congruence of the signals and the direction of the visuo-vestibular input on these vestibular regions.

**Table 2.** Coordinates used to generate the regions of interest (ROI) regarding the second ROI analysis on the visuo-vestibular regions.

| # | label | left coordinates |  |  | right coordinates |  |  |
| --- | --- | --- | --- | --- | --- | --- | --- |
|  |  | x | y | z | X | y | z |
| 1 | V6 | -13 | -86 | 28 | 15 | -80 | 31 |
| 2 | VIP | -45 | -43 | 4 | 40 | -51 | 43 |
| 3 | CSv | -11 | -27 | 43 | 11 | -29 | 44 |
| 4 | MT | -46 | -63 | 0 | 43 | -63 | 0 |
| 5 | PIC | -42 | -21 | 21 | 43 | -23 | 18 |

Coordinates were converted from Talairach [1] to MNI space (see Methods section). V6 = visual area 6, VIP = ventral intraparietal area, CSv = cingulate sulcus visual area, hMT+ = human middle temporal visual area, PIC = posterior insular cortex.

## Results

### Visuo-vestibular multimodality network

This study first aims at localizing the cortical regions with increased responses to self-motion signals when vestibular input is concurrent to unambiguous optic flow patterns. To investigate this issue, the brain activations produced when stimulating both the vestibular and visual systems were contrasted to those induced when recruiting only the visual modality. The results of this contrast are shown on the inflated left and right cortical surfaces (Fig. 2A) and on horizontal sections (Fig. 2B) of template brains. It can be seen that the activations specific to the simulataneous visuo-vestibular stimulation are concentrated in the anterior and posterior insular regions, the lateral sulcus, the temporo-parietal junction, the superior temporal gyrus and the inferior parietal lobule. In total, we found seven bilateral activation sites whose coordinates are given in Table 1, together with the corresponding second-level t-values. By using the Anatomy3 toolbox [30], the Brainnetome atlas [33] and a review of the literature regarding the vestibular responsive brain regions, we propose that region 1, which lies in the ventral granular insula, might correspond to the anterior insula (AI) region [9, 25, 34, 42, 67]. Region 2 is located in the dorsal granular insula and might correspond to area operculum 3 (OP3) [30, 31, 42]. Region 3, located in the frontal opercular cortex, could match with area operculum 8 (OP8) [4, 30]. A second region located in the dorsal granular insula, region 4, might correspond to previously established coordinates [36, 42, 67] of the parieto-insular vestibular cortex (PIVC) [38, 39, 41, 46]. It tends to show a geographical overlap with area operculum 2 (OP2) [31, 42], often referred to as a major hub in the human vestibular network [32, 35, 36], activated by vestibular inputs but not by visual motion [19, 36]. Region 5, which lies in the central opercular cortex, might correspond to area operculum 4 (OP4) [30, 42]. By the temporo-parietal junction (tpj), located at the lower end of the postcentral gyrus, region 6 coordinates find correspondence with previous studies [35, 36, 67] identifying this region as the posterior insular cortex (PIC) [35, 42]. Finally, region 7, located between the superior temporal gyrus (stg) and the inferior parietal lobule (ipl) in the planum temporale gyrus, might correspond to PFcm (area supramarginalis magnocellularis columnata), one of the seven cytoarchitectonically defined “inferior parietal lobule (IPL) areas” [18, 42]. Note that the opposite contrast (“Visual only” > “Visual + vestibular”) reveals only one bilateral region located close to the central sulcus (MNI coordinates [46-10 34] and [-42-10 34]), nearby both the somaesthetic or motor cortices and it corresponds to a region previously described in the literature as bilaterally deactivated when the vestibular saccule is stimulated [56].

**Fig. 2.**
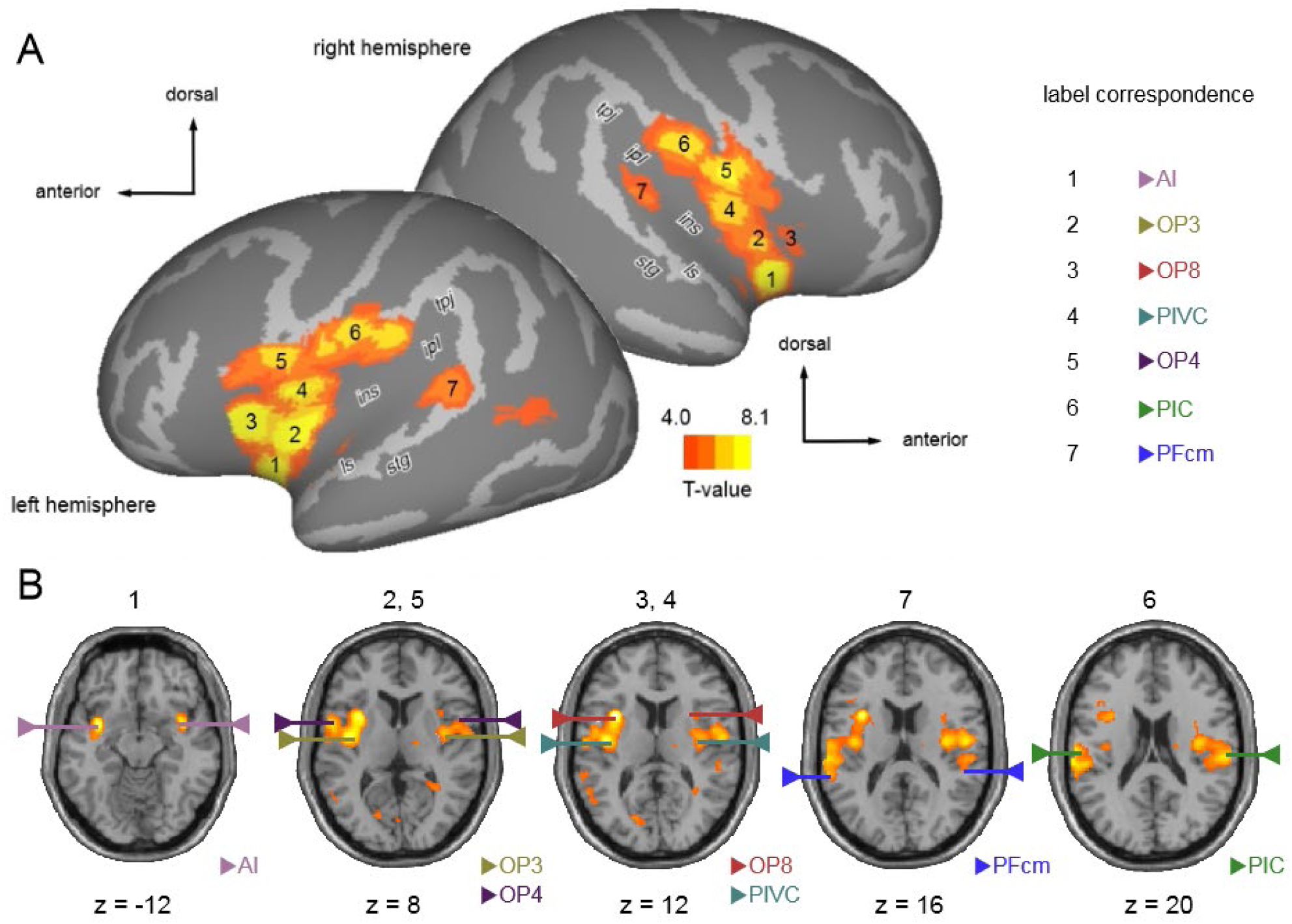
Brain activations for the contrast “Visual + vestibular” > “Visual only”. A/ Statistical parametric map for the contrast “Visual + vestibular” > “Visual only” (second level analysis; n = 15 participants) projected on an inflated brain template (A) and on horizontal slices of the SPM12 single subject T1 template (B). Seven regions exhibited statistical significance (t-value > 4, p < 10^-3^ uncorrected) in both left and right hemispheres. Coordinates and t values of their local maxima are provided in Table 1. (Anatomical landmarks: ls = lateral sulcus, ins = insula, stg = superior temporal gyrus, ipl = inferior parietal lobule, tpj = temporoparietal junction). The number(s) above the slice in (B) indicates the number of the region(s) represented. The ends of the lines designate the location of the area and the name of the region is indicated in the corresponding color in the bottom right-hand corner of each slice.

### Activation preferences of regions solicited by visuo-vestibular stimulation

To further investigate whether the above described regions exhibit more specific functions related to the visuo-vestibular integration process, we performed ROI analyses to assess (1) their change in activation when the stimulation is concurrently visual and vestibular compared to visual only, (2) their preference for congruent visuo-vestibular signals, and (3) their specificity to forward self-motion as specified by these signals. Fig. 3 shows the location of the spherical ROIs on lateral and medial views of the template’s inflated cortical surfaces (Fig. 3A), together with the results of the ROI analyses for the “Visual + vestibular” > “Visual only” contrast (Fig. 3B), the “Congruent” > “Incongruent” contrast (Fig. 3C) and the “Forward” > “Backward” contrast (Fig. 3D). Note that (1) since the activation patterns were visually very similar on both hemispheres (see Fig. 2A and Fig. 2B), left and right ROIs were grouped together to increase statistical power (n = 30 hemispheres), and (2) PFcm (region 7) was excluded from the present analyses because its statistical significance did not hold when using only half of the 15 subjects’ runs. The “Visual + vestibular” > “Visual only” contrast (Fig. 3B) serves here only as a positive control for our statistical analysis and as a point of comparison with the second ROI analysis. As expected, since this is the contrast on which they had been identified, all the regions of interest (AI, OP3, OP4, OP8, PIC, PIVC) show highly significant mean contrast values (***, p < 0.001). Among the above mentioned regons, none of them showed significant activation, neither for the “Congruent” > “Incongruent” contrast (Fig. 3C), nor for the “Forward” > “Backward” contrast (Fig. 3D).

**Fig. 3.**
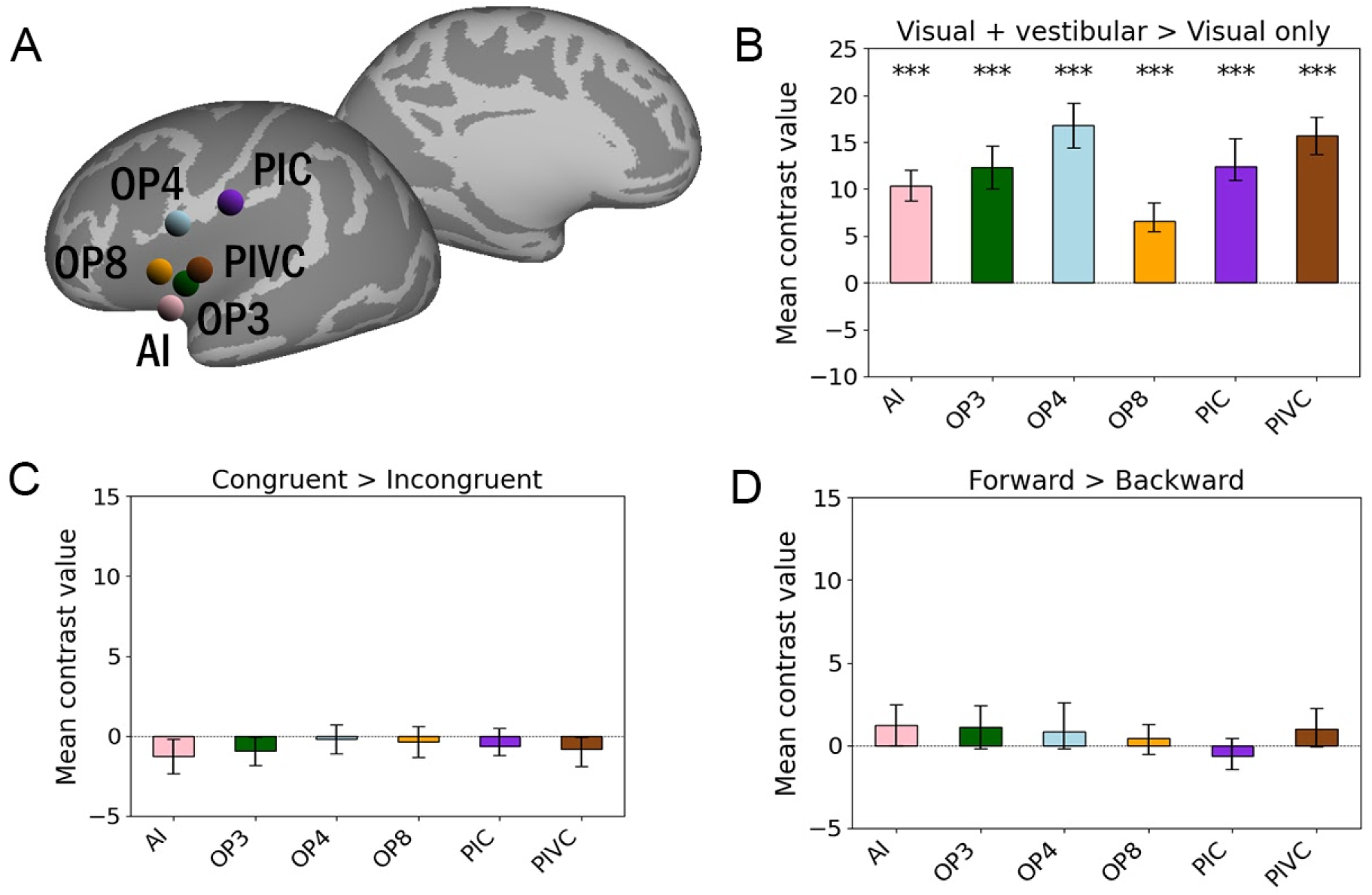
ROI analysis for the regions defined with the “Visual + vestibular” > “Visual only” contrast. Mean contrast value within each ROI is the average of that ROI in all hemispheres (n = 30). A/ ROIs’ location on the lateral and medial views of an inflated brain template. Bar plot of mean contrast values in each ROI regarding the “Visual + vestibular” > “Visual only” contrast (B), the “Congruent” > “Incongruent” contrast (C) and the “Forward” > “Backward” contrast (D). In all bar plots, error bars indicate the 90% confidence interval (IC90) across hemispheres (n = 30). One-sample one-tail t-tests with the hypothesis that the values are strictly superior to zero were used to assess the significance of each ROI (* p < 0.05, ** p < 0.01, *** p < 0.001).

### Activation preferences of the regions involved in the processing of anteroposterior vestibular input

Many brain regions known to process both visual and vestibular input [14] were not activated by our main contrast (“Visual + vestibular” > “Visual only”, see Fig. 2A and Fig. 2B). We postulate that with a relatively high motion-coherence (30%), the unambiguous and full contrast nature of our optic flow stimuli was sufficient to evoke maximal activations in those regions and that the addition of a simultaneous vestibular input could not further increase the activation level. Nevertheless, we wanted to know whether some of these regions could actually exhibit specific functions related to the visuo-vestibular integration process, by contrast with the vestibular-dominant regions described above. Building on our previous study [1], in which optic flow and GVS were administered separately, we constructed ROIs centered in the coordinates given in that previous work for V6, VIP, the cingulate sulcus visual area (CSv), the human middle temporal visual area (hMT+) and the posterior insular complex (PIC) (see Table 2 for their MNI coordinates). Fig. 4 shows the location of the spherical ROIs on the template’s inflated cortical surfaces (lateral and medial view) (Fig. 4A) together with the results of the ROI analyses for the “Visual + vestibular” > “Visual only” contrast (Fig. 4B), the “Congruent” > “Incongruent” contrast (Fig. 4C) and the “Forward” > “Backward” contrast (Fig. 4D), with the same conventions as those used in Figure 3. Regarding the “Visual + vestibular” > “Visual only” contrast, although PIC showed a highly significant activation (***, p < 0.001), as expected from the previous ROI analysis, area V6 was also found to exhibit very significant activation (**, p < 0.01), by contrast with all the other regions considered in this second ROI analysis. In the case of the “Congruent” > “Incongruent” contrast, a single region, V6 again, shows a very significant difference (**, p < 0.01) between the two conditions, revealing a preference for congruent signals. Finally, in the case of the “Forward” >” Backward” contrast, the same single region, V6, shows a highly significant difference (***, p < 0.001), indicating that area V6 prefers signals specifying forward self-motion. Therefore, within the 5 regions considered in this second ROI analysis (V6, VIP, CSv, hMT+, PIC) only V6 shows preferential activation when when concurrent visual and vestibular signals are congruent and when the direction of the self-motion information is in the forward direction.

**Fig. 4.**
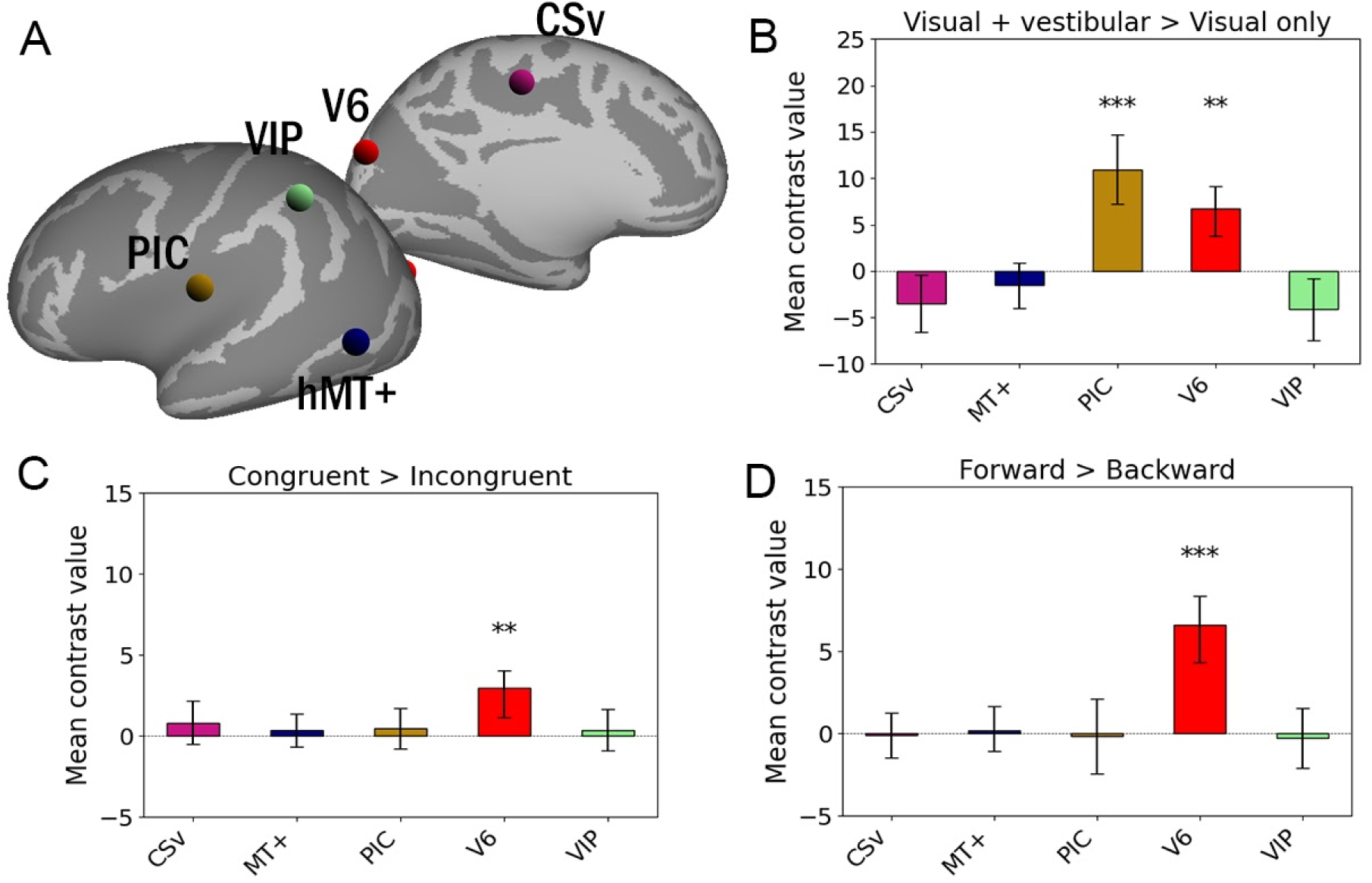
ROI analysis for the regions initially defined in our princeps study [1] (see Table 2). Mean contrast value within each ROI is the average of that ROI in all hemispheres (n = 30). A/ ROIs’ location on an inflated brain template (lateral and medial views). Bar plot of mean contrast values regarding the “Visual + vestibular” > “Visual only” contrast (B), the “Congruent” > “Incongruent” contrast (C) and the “Forward” > “Backward” contrast (D). In all bar plots, error bars indicate the IC90 across hemispheres (n = 30). One-sample one-tail t-tests with the hypothesis that the values are strictly superior to zero were used to assess the significance of each ROI (* p < 0.05, ** p < 0.01, *** p < 0.001).

## Discussion

This study aimed to more precisely characterize the brain regions involved in integrating visual and vestibular signals, a necessary step for the perception and control of self-motion. We also addressed their possible specificities regarding signal congruence and direction of self-motion cue. Our results revealed a consistent bilateral network that is particularly engaged when the stimulation is both visual and vestibular. This network is composed of cortical areas (PIVC, PIC, OP3, OP4, PFcm and AI) previously proposed to be parts of the multimodal vestibular cortex [42] and a lesser-known region, OP8, whose role in multisensory integration or self-motion processing had not been reported so far. These results reinforce and complete the current knowledge regarding the mapping of the network involved in the multimodal processing of sensory information linked to the perception and/or control of self-motion [20, 54]. Our subsequent ROI analysis indicate that none of the above-mentioned areas show any differential activity either between congruent and incongruent visuo-vestibular information, or between forward and backward self-motion cues (Fig. 3). Importantly, these results do not rule out the possibility that underlying neurons can actually be sensitive to these factors but simply that their overall activity does not reflect any global preference for congruence of forward self-motion. We extended our study by repeating these ROI analyses on other regions known to be involved in the processing of both visual and vestibular information. The regions concerned by this second ROI analysis are V6, VIP, CSv, MT+ and PIC [1] (see Table 2). When the stimulation is visuo-vestibular and not solely visual (Fig. 4B), two of these five regions, V6 and PIC, show significantly greater activation, indicating that they are the most influenced by the concurrent presence of visual and vestibular input. Concerning the discrimination of congruent and incongruent conditions (Fig. 4C), only one region, V6 again, shows significantly more activation in the “Congruent” conditions compared to the “Incongruent” conditions. V6. Therefore, V6 seems to be specialized in detecting the congruence of visuo-vestibular signals, a function previously unknown to this region. To our knowledge, the only previous study examining the cortical response to congruence or incongruence of visual and vestibular signals in humans was conducted by Roberts *et al.* [53], which found preferential activation of the PIVC region in the case of signal incongruence. However, the aforementioned study focused on movement input towards the left or the right, not in the anteroposterior direction, and used caloric vestibular stimulation [47] rather than GVS, suggesting that our new results are not contradictory but complementary. Last but not least, regarding the discrimination between forward and backward self-motion, the V6 area is again the only region to exhibit a significantly greater activation when the visuo-vestibular stimulation mimics forward self-motion (Fig. 4D). Therefore, this region appears to be even more specialized than we initially thought [1], showing a marked preference not only for the anteroposterior axis, but for the forward direction.

### Visuo-vestibular signals integration

The processing of visual stimulation consistent with self-motion is known to engage, in humans, areas such as V6, VIP, CSv, PIC and MT+, which have also been shown to be activated by vestibular galvanic stimulation [1]. However, it is worth noting that when we analyze the “Visual + vestibular” > “Visual alone” contrast, out of the mentioned regions, PIC, PIVC and V6 are the only regions that appear to respond more strongly when the stimulation is visuo-vestibular and not only visual (see Fig. 3 and Fig. 4). As only signals of motion in the anteroposterior axis were probed, it cannot be ruled out that some regions within the visuo-vestibular cortical network are not sensitive to self-motion cues in the anteroposterior axis, but are instead dedicated to other axes. Hence, PIC, PIVC and V6 are three sites of visuo-vestibular integration sensitive to the presence of concurrent vestibular information, even when the visual signal is already sharp. As for the other regions identified as visuo-vestibular but not observed on our contrast, we suggest that they are less sensitive to the presence of a simultaneous vestibular input and a sharp visual signal is sufficient to saturate their activation. In other multisensory stimuli, such as visual-auditory stimuli, it has been shown that compensation between the two modalities was most apparent when one of the stimuli was ambiguous [66]. Consequently, it is likely that under conditions where visual stimulation compatible with self-motion would be less sharp, vestibular compensation would be more readily identifiable. Recent psychometric studies have established that a clear vestibular signal can resolve ambiguous visual motion [3], and that the visuo-vetsibular integration could follow a weighted-averaging rule that depends upon the relative reliability of each modality [45].

To our knowledge, similar devices have not yet been implemented in the context of MRI acquisitions to identify the cortical correlates of this phenomenon. Thus, generating optic flow stimuli with a lower percentage of motion-coherence might provide better visualization of cortical activation in response to vestibular modality intervention. We believe that it would be useful in future protocols to include one or more conditions (depending on the number of axes and directions selected) involving only the vestibular modality. This additional condition would complement the proposed contrasts and would enable the use of tools as the multisensory enhancement index, “which compares the bimodal response to the largest unimodal response” and the additivity index, “which compares the bimodal response to the sum of the unimodal responses” [24, 63]. It should be noted that the present study investigates the integration of simultaneous visual and vestibular signals at the cortical level, deliberately without reporting any perceptual aspect of the subject. Our aim here was to decipher the mechanisms involved in integration following simultaneous stimulation of the visual and vestibular organs, and not in response to the subject’s own perception of self-motion. The cortical response to the conscious perception of simultaneous visual and vestibular stimuli could be studied in a similar way to ours, using longer and/or more intense stimuli, but deserves further research with an adapted protocol [54].

### V6 involvement in self-motion signals integration

Our results reveal a great and unique specificity of area V6 in the processing of self-motion information in the anteroposterior axis, particularly in the context of congruent visuo-vestibular signals and in the forward direction. While the evolution of the human brain and visuospatial integration is also a subject of study in the field of anthropology [14], it is not surprising to imagine that cortical regions such as V6 are particularly specialized in the processing of information related to self-motion in the most ecological situation for humans (*i.e.,* forward movement with visuo-vestibular stimulation). Through visual testing paradigms of self-motion perception, V6 was quickly recognized as an important region in the integration of sensory information useful for locomotion [8, 64], responding well to optic flow patterns while also showing a sensitivity to stereoscopic depth gradients [16, 17]. Subsequent studies further documented its involvement in the multisensory integration of signals linked to self-motion, with vestibular [1, 15] and proprioceptive sensitivity [55]. V6 is undoubtedly a key region for the multisensory integration of information necessary for locomotion. Within the present study, we make an important contribution to the understanding of its role by highlighting, for the first time, a specialization for ecologically-meaningful self-motion signals; that is, signals that are congruent across modalities and specify the most common —forward— direction of self-motion.

## Conclusion

In this study, we document a consistent bilateral cortical network with increased activity when vestibular self-motion signals coincide with visual ones. Importantly, we also demonstrate that area V6 plays a pivotal and unique role in visuo-vestibular integration within the context of ecologically relevant forward self-motion. These findings provide new insights into our understanding of cortical self-motion processing and multisensory integration and may help in comprehending the pathologies associated with deficient visuo-vestibular integration.

## ACKNOWLEDGEMENTS

We thank the INSERM U1214 MRI technical platform for the MRI acquisitions. The fMRI recordings were funded by a grant from the Institut des Sciences du Cerveau de Toulouse (ISCT). We would also like to thank the Agence Nationale de la Recherche (ANR) for funding the In-Vest projet (ANR-21-CE37-0023-01) and Brandon Hayes, Jean-Pierre Jaffrézou and Simona Celebrini for their thorough revision of English grammar and spelling.

